# BRG1 is a prognostic indicator and a potential therapeutic target for prostate cancer

**DOI:** 10.1101/506972

**Authors:** Rohini Muthuswami, LeeAnn Bailey, Radhakrishnan Rakesh, Anthony N. Imbalzano, Jeffrey A. Nickerson, Joel W. Hockensmith

## Abstract

BRG1 is one of two mutually exclusive ATPases that function as the catalytic subunit of human SWI/SNF chromatin remodeling enzymes. BRG1 has been identified as a tumor suppressor in some cancer types but has been shown to be expressed at elevated levels, relative to normal tissue, in other cancers. Using the TCGA (The Cancer Genome Atlas) prostate cancer database, we determined that BRG1 mRNA and protein expression is elevated in prostate tumors relative to normal prostate tissue. Only 3 of 491 (0.6%) sequenced tumors showed amplification of the locus or mutation in the protein coding sequence, arguing against the idea that elevated expression due to amplification or expression of a mutant BRG1 protein is associated with prostate cancer. Kaplan-Meier survival curves showed that BRG1 expression in prostate tumors inversely correlated with survival. However, BRG1 expression did not correlate with Gleason score/ISUP Grade Group, indicating it is an independent predictor of tumor progression/patient outcome. To experimentally assess BRG1 as a possible therapeutic target, we treated prostate cancer cells with a biologic inhibitor called ADAADi that targets the activity of the SNF2 family of ATPases in biochemical assays but showed specificity for BRG1 in prior tissue culture experiments. The inhibitor decreased prostate cancer cell proliferation and induced apoptosis. When directly injected into xenografts established by injection of prostate cancer cells in mouse flanks, the inhibitor decreased tumor growth and increased survival. These results indicate the efficacy of pursuing BRG1 as both an indicator of patient outcome and as a therapeutic target.

## Introduction

Prostate cancer is the most common cancer in men, is the second leading cause of cancer death in men in the United States, and is the fifth leading cause of cancer death in men worldwide (http://globocan.iarc.fr/old/FactSheets/cancers/prostate-new.asp; https://www.cancer.org/cancer/prostate-cancer.html). Though treatable, definitive understanding about the molecular origins of prostate cancer is lacking, and there is considerable controversy about screening methods (1–6). Continued efforts to identify molecular markers that distinguish between tumors that will remain latent, progress slowly, or progress aggressively are needed (7,8).

There are two closely related, mutually exclusive ATPases that function as the catalytic subunits of the human/mammalian SWI/SNF chromatin remodeling enzymes, termed BRG1 and BRM (9–13). These enzymes are widely utilized in the cell to regulate gene expression, replication, repair, recombination, and higher–order genome organization. Not surprisingly, both enzymes have been implicated in diverse types of cancer (14–16). There is considerable evidence that both the BRG1 and the BRM enzymes are mutated or exhibit altered expression in many cancers, but to date, the evidence suggests that the consequences of these changes vary widely depending on the type of cancer.

Loss of BRG1 function has been shown in a number of cancers, most notably small cell carcinoma of the ovary, hypercalcemic type (17–20), and non-small cell lung cancers (21–23), and results in loss of a number of cell functions related to tumor suppression. BRM loss has been implicated in a number of tumor types, and the idea of targeting BRM in BRG1-deficient cancers has received attention as a potential therapeutic strategy (24–26). In contrast, there is emerging evidence that BRG1 is expressed at elevated levels in some tumors relative to normal tissue. This includes disparate tumor types such as breast cancer, melanoma, neuroblastoma, colorectal cancer, and prostate cancer (27–33). Though increased expression of mutated proteins could be consistent with a tumor suppressive function, there is no evidence of prevalent BRG1 or BRM mutation in breast cancer or melanoma (34,35), where such analysis has been reported. Elevated BRG1 expression has been linked to numerous pathways converging on cell proliferation and survival, including SHH and WNT signaling, the PI3K/AKT pathway, and regulation of lipogenesis and ABC transporter induction (reviewed in (16)).

These results indicate that, if properly delivered, inhibitors of BRG1 function may represent a potential therapeutic approach to certain cancers. PFI-3, a small molecule bromodomain inhibitor with structural specificity for three bromodomain containing subunits of the mammalian SWI/SNF enzymes (BRG1, BRM, and polybromo-1, also called BAF180) (36) has been shown to modulate certain cell differentiation transitions (37,38) but has no effect whatsoever on cancer cell proliferation (36,39). However, promising results for inhibition of cancer cell proliferation have resulted from studies of an as yet structurally undefined inhibitor of the SNF2 family of ATPases called ADAADi (Active DNA-dependent ATPase A Domain inhibitor). ADAADi is a chromatographically separable byproduct of the bacterial aminoglycoside-phosphotransferase (APH) action upon aminoglycosides (40,41). ADAADi has largely been used as a biochemical probe to help define enzymatic activities of SNF2 ATPases in vitro (40–42), and it therefore functions as an inhibitor of other ATP-dependent activities of these enzymes, such as chromatin remodeling (41). In tissue culture, ADDADi inhibits cancer cell proliferation and survival (40,43) and there appears to be some specificity of ADDADi for BRG1, as both ADAADi and shRNA-mediated knockdown of BRG1 inhibited proliferation, but there was no additive effect of the inhibitor plus BRG1 knockdown (43). This result suggests that even if other related ATPases are contributing to cell proliferation and survival, their contributions are relatively minor. In addition to effects on cancer cell proliferation, ADDADi phenocopies BRG1 knockdown in demonstrating essential functional roles for BRG1 in cancer cell metabolism and in drug-induced activation of ABC transporter proteins linked to chemoresistance (39,43). A specific BRG1-targeting molecule may therefore be of therapeutic value in treatment of certain cancers.

In this report, we interrogated the TCGA Prostate Cancer database for links between human prostate tumors and BRG1 expression. We found that BRG1 expression, as well as the expression of the related ATPase, BRM, is elevated in prostate cancer biopsies relative to normal prostate tissue. The mutation rate for BRG1 and BRM in prostate cancer is less than 1% for each protein. Stratifying tumor samples by expression level revealed an inverse relationship between BRG1 expression and patient outcome, indicating BRG1 is a prognostic indicator for prostate cancer. Interestingly, there was no correlation between BRG1 expression and the Gleason score of the tumor samples, indicating that BRG1 is a prognostic indicator that is independent of the microscopic features of the tumor. Challenging prostate cancer cells in tissue culture with ADAADi led to decreased proliferation and survival, with evidence of apoptosis among the dying cells. When prostate cancer cells were used to orthotopically seed tumors in mouse flanks, direct injection of ADAADi, but not other compounds, inhibited tumor growth and improved mouse survival.

## Materials and methods

### Analysis of TCGA Prostate Cancer Patient Data

Patient data was from The Cancer Genome Atlas (TCGA) prostate adenocarcinoma (PRAD) dataset (35). Almost all prostate cancer is adenocarcinoma. The last update for the TCGA PRAD data is 5/31/2016 and includes data from 498 tumors and 52 normals. Various online programs were used to analyze this data. TCGA PRAD expression data was analyzed using Xena, a functional genomics and analysis platform developed by the University of California at Santa Cruz (xena.ucsc.edu) (44). Correlations between BRG1 and BRM mRNA levels in patient tumors were plotted using GEPIA (Gene Expression Profiling Interactive Analysis), a web-based tool for analysis of TCGA datasets (gepia.cancer-pku.cn) (45). GEPIA was also used to draw boxplots and to correlate disease free survival (DFS, also called relapse-free survival and RFS) with gene expression. GEPIA uses a Log-rank test, or the Mantel–Cox test, for survival hypothesis testing. The Cox proportional hazard ratio and the 95% confidence interval information are also calculated. BRG1 and BRM mutations in TCGA prostate tumors were analyzed by CBioPortal (CBioPortal.org) (46,47). Correlations between Gleason Scores and gene expression were done using Betastasis (http://www.betastasis.com/prostate_cancer/tcga_prad_from_gdc).

#### Synthesis and purification of ADAADi

ADAADi for cell culture studies was synthesized and purified as described (40). ADAADi for animal studies was synthesized and purified as described (41).

#### Cell lines

Human prostate adenocarcinoma cell line PC3 was purchased from ATCC or from NCCS, Pune, India and maintained in Dulbecco‘s Modified Eagle Medium (DMEM) supplemented with 10% fetal bovine serum (FBS), 1% penicillin-streptomycin-amphotericin cocktail at 37°C in the presence of 5% CO_2_.

#### Cell viability assay

5000 cells were seeded in each well of a 96-well plate containing 200 μl media and incubated overnight. The media was replaced with media containing varying concentrations of ADAADi. The cells were incubated at 37°C for the indicated time point. The media in each well was then replaced with media containing 0.45 mg/ml MTT and incubated for 2 hours at 37°C in a CO_2_ incubator. The media containing MTT solution was discarded and the purple precipitate formed was dissolved in 100 μl isopropanol. The plate was incubated for 15 min at room temperature and the absorbance was measured at 570 nm.

#### Annexin V-FITC apoptosis detection

PC3 cells were grown in 60 mm dish to 60-70% confluency before treatment with sub-lethal concentration (5 μM) of ADAADi for 48 hr. The cells were stained with Annexin V and PI using Annexin V-FITC apoptosis detection kit (eBiosciences, USA; Catalog # BMS500FI/100). The samples were analyzed using BD FACS Calibur 4C flow cytometer.

### Mouse xenograft studies

All animal experiments were performed according to protocols approved by the Institutional Animal Care and Use Committee at the University of Virginia School of Medicine. CRL: CD-1nu/nu Br mice were obtained from Charles River Animal Resources Facility. PC3 cells were grown in RPMI 1640 media containing 10% serum. 200 μl of this cell culture was diluted 1:1 with Matrigel (Collaborative Biomedical Products) such that the total number of cells after dilution were ~2 × 10^6^. The cells were then injected on the underside of the flank and the development of subcutaneous tumor size was monitored by measurement with Vernier calipers.

Control and ADAADi treatments were started when the tumor size reached 200 mm^3^. ADAADi was solubilized in PBS and pH was adjusted to 7.2 using phosphoric acid and filter sterilized before injection. 50 μl of the drug was administered by direct injection into the tumor. In two independent experiments, injections were repeated every other day for two weeks. Similarly, a third experiment utilized the every other day protocol for two weeks and after a one week break from treatment, an additional set of injections were performed, again, every other day for two weeks. The mice were euthanized when the tumor size exceeded 1000 mm^3^ or when the weight of the mice decreased by more than 15% of their starting weight. The tumor size was calculated using the following formula: volume = [length x (width)^2^] / 2.

## Results

### BRG1 expression is elevated in prostate tumors relative to normal tissue

BRG1 and BRM, two closely related ATPases that are mutually exclusive catalytic subunits of mammalian SWI/SNF chromatin remodeling enzymes, have differing functional roles in different types of cancer (14–16). Prior reports about BRG1/BRM in prostate cancer have utilized immunohistochemistry (IHC) of patient tumor samples and determined that BRG1 protein levels were higher in tumor than in normal tissue. BRM protein levels were described as heterogeneous, with an average value suggesting BRM expression in tumors was lower than in normal tissue. cDNA microarray analysis was generally consistent with the evaluation of protein expression by IHC (48,49). We took a complementary approach to evaluating BRG1/BRM expression in prostate cancer by interrogating the Cancer Genome Atlas (TCGA) database. Data from 45 patients with matched normal tissue and prostate tumors were evaluated. BRG1 expression, on average, was significantly elevated in adenomas relative to normal tissue, while average BRM mRNA expression was significantly decreased in tumor relative to normal tissue (Table 1). As controls for our analyses, we evaluated prostate serum antigen (PSA; KLK3), which showed significant elevation in expression in tumors, and SUN1, an inner nuclear envelope protein not previously linked to prostate cancer, which showed no significant difference (Table 1). Boxplots showing the ranges for BRG1 and BRM mRNA expression in all TCGA samples (n = 492 for tumors, n= 52 for normal) reinforced the conclusion that BRG1 mRNA expression is elevated in tumor samples compared to normal tissue while the converse is true for BRM mRNA (Fig. 1A). The UCSC Xena tool was used to determine whether any correlation between BRG1 and BRM mRNA expression in prostate cancer exists; the results clearly show no correlation across the dataset (Fig. 1B). To evaluate protein expression, we queried published proteomic data for BRG1 from 28 prostate tumor and 8 normal prostate tissue samples (50). We were unable to identify similar proteomic data for BRM. The data show a statistically significant increase in BRG1 protein expression in tumor relative to normal tissue (Table 2) that matches the magnitude of the increase in BRG1 mRNA determined from analyzing the TCGA dataset (Table 1, Fig. 1A).

**Figure 1:**
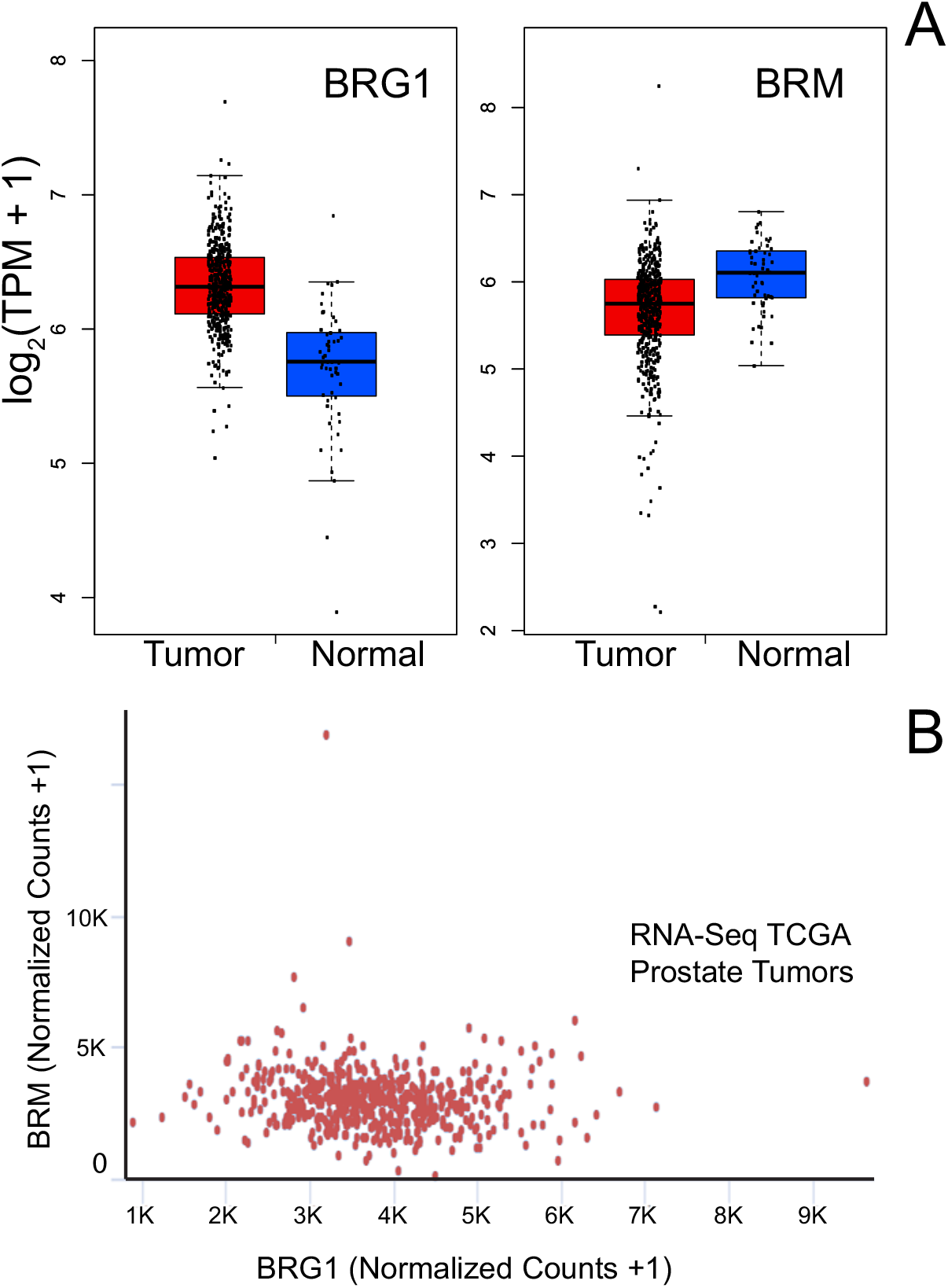
BRG1 mRNA expression in prostate cancer patient tissue is elevated compared to that in normal prostate tissue while BRM mRNA expression in prostate cancer patient tissue is reduced compared to that in normal prostate tissue. All data was extracted from the prostate adenocarcinoma TCGA dataset. Plots were generated with GEPIA software (45). **(A)** Boxplots for BRG1 (left) and BRM (right) mRNA expression in prostate tumors (red) compared to normal prostate tissue (blue). Boxes enclose the middle two quartiles of mRNA expression with a centerline at the median. **(B)** BRG1 and BRM mRNA levels are not correlated in prostate tumors.

**TABLE 1:**
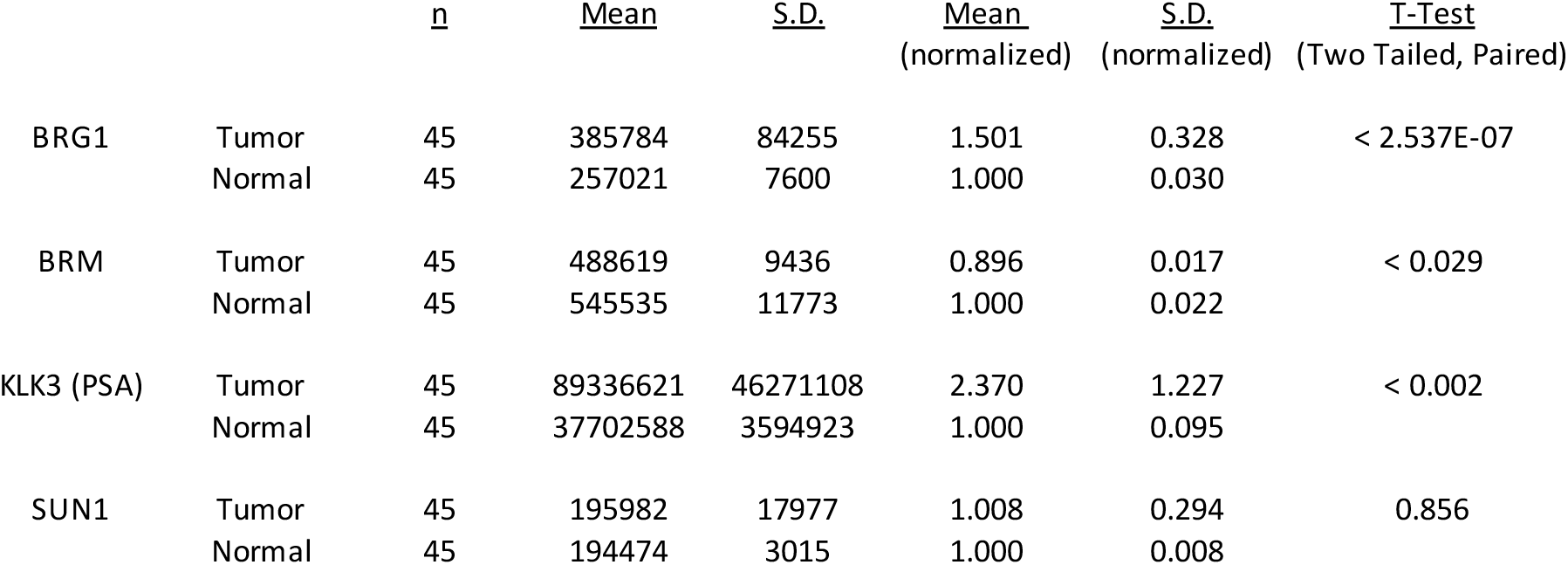
mRNA expression in matched prostate tumor and normal tissue samples from TCGA

|  |  | <u>n</u> | <u>Mean</u> | <u>S.D.</u> | <u>Mean</u><br>(normalized) | <u>S.D.</u><br>(normalized) | <u>T-Test</u><br>(Two Tailed, Paired) |
| --- | --- | --- | --- | --- | --- | --- | --- |
| BRG1 | Tumor | 45 | 385784 | 84255 | 1.501 | 0.328 | < 2.537E-07 |
|  | Normal | 45 | 257021 | 7600 | 1.000 | 0.030 |  |
| BRM | Tumor | 45 | 488619 | 9436 | 0.896 | 0.017 | < 0.029 |
|  | Normal | 45 | 545535 | 11773 | 1.000 | 0.022 |  |
| KLK3 (PSA) | Tumor | 45 | 89336621 | 46271108 | 2.370 | 1.227 | < 0.002 |
|  | Normal | 45 | 37702588 | 3594923 | 1.000 | 0.095 |  |
| SUN1 | Tumor | 45 | 195982 | 17977 | 1.008 | 0.294 | 0.856 |
|  | Normal | 45 | 194474 | 3015 | 1.000 | 0.008 |  |

**TABLE 2:** BRG1 protein levels in prostate tumor and normal tissue samples extracted from ref. 50

|  | <u>n</u> | <u>Mean</u> | <u>S.D.</u> | <u>Mean</u><br>(normalized) | <u>T-Test</u><br>(Two Tailed, Paired) |
| --- | --- | --- | --- | --- | --- |
| normal | 8 | 0.716 | 0.180 | 1.000 |  |
| tumor | 28 | 1.054 | 0.583 | 1.473 | < 0.013 |

We examined sequence data for the BRG1 and BRM coding sequences to determine whether any of the patient tumors were mutated for either or both ATPases. Only two of 491 patients had BRG1 mutations: one with a V698I and G775D double substitution and one with an I1214L substitution (Fig. 2A). In addition, one patient showed amplification of the BRG1 locus. No patients with deletions in the BRG1 coding sequence were identified. Three patients had BRM mutations, one had amplification of the locus, and ten patients with homodeletions were identified (Fig. 2B). One BRM mutation was an R1159Q substitution; another was a frameshift after amino acid 1253. Interestingly, the patient with the double substitution in BRG1 also had a R524M substitution in BRM. The significance of the mutation in both proteins is unknown. Regardless, the data indicate that a small percentage of prostate tumors contain mutations in either BRG1 or BRM. Thus the possibility that elevated expression of mutant BRG1 or BRM protein correlates with the prostate tumor phenotype is not supported.

**Figure 2:**
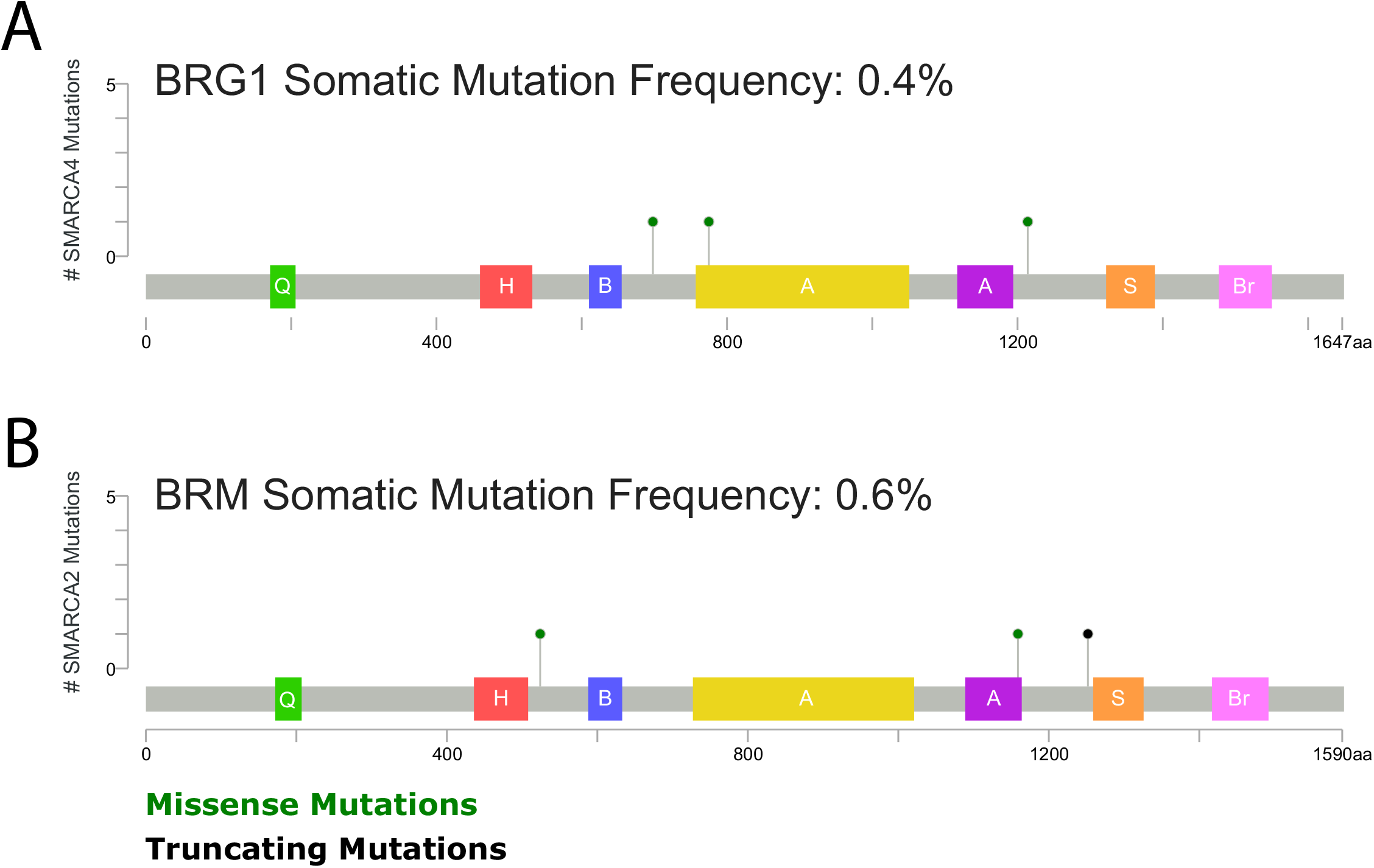
Schematic diagram of the location of BRG1 and BRM somatic mutations present in TCGA prostate tumor patients relative to known domains of BRG1 and BRM proteins. CBioPortal (46,47) was used for data analysis. Domains indicated: Q (green) – QLQ domain, H (red) – HSA domain, B (blue) – BRK domain, A (yellow and purple) – bipartite SNF2 ATPase domain, S (orange) – SnAC domain, Br (pink) – bromodomain.

### BRG1 is a prognostic indicator for prostate cancer patient outcome

Despite the links between increased BRG1 expression and prostate tumors, there is presently no understanding of whether BRG1 expression correlates with patient outcome. We stratified patient BRG1 mRNA expression data from the TCGA database and compared patient survival among those in the highest quartile of BRG1 mRNA expression and those in the lowest quartile. Kaplan-Meier plots demonstrate a significant difference in the two patient populations, with BRG1 expression inversely correlating with survival (Fig. 3A). In contrast, similar analysis of BRM expression in prostate cancer patients showed no correlation between BRM expression and patient outcome (Fig. 3B). The results indicate that BRG1 is a prognostic indicator of prostate cancer patient outcome.

**Figure 3:**
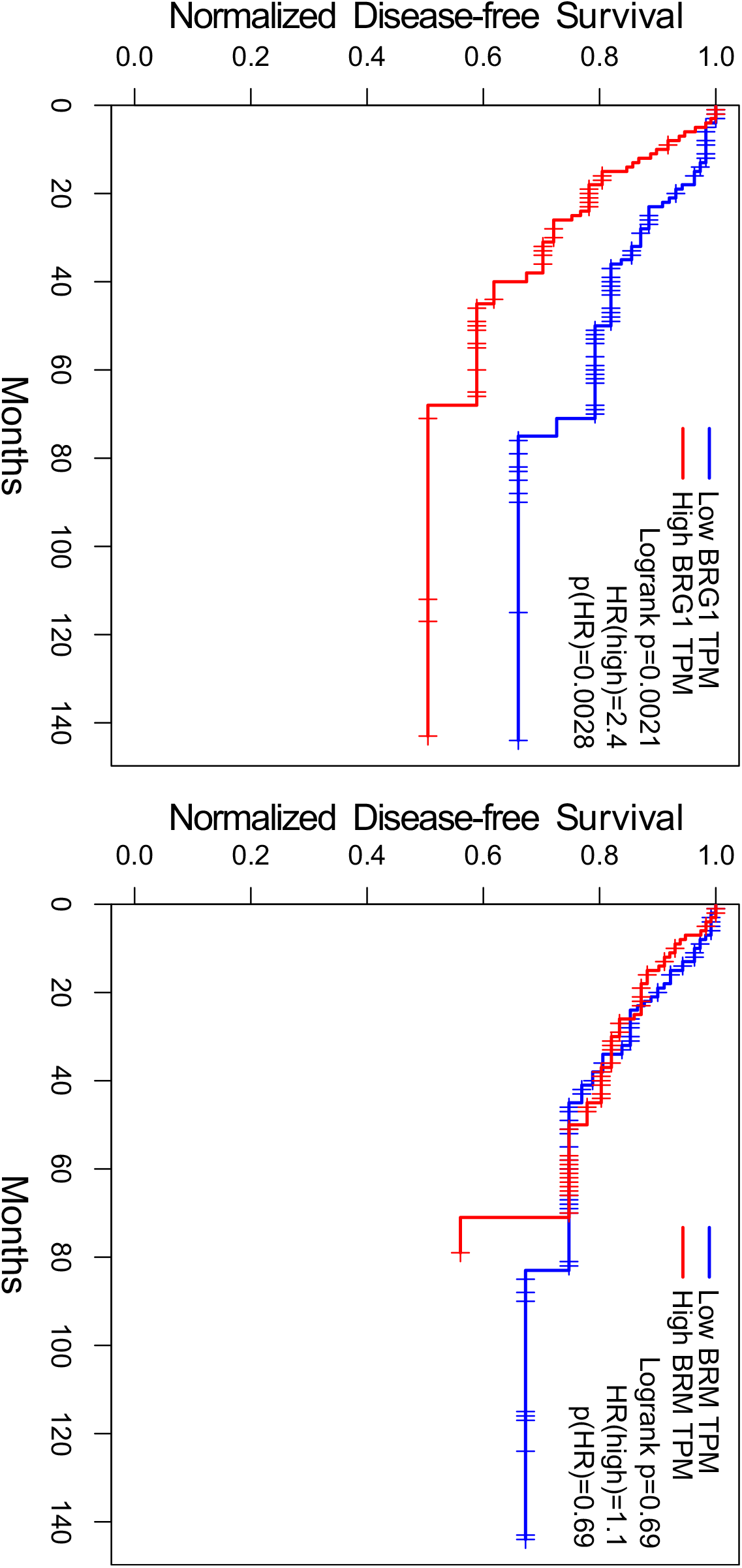
BRG1 mRNA levels **(A)** but not BRM mRNA levels **(B)** inversely correlate with prostate tumor patient survival. Kaplan-Meier plots shown are based on analysis of TCGA prostate patient data using GEPIA (45). The blue line labelled “Low” is the patients with the lowest quartile of mRNA levels; the red line labelled “High” is the patients with the highest quartile of mRNA levels. TPM, transcripts per kilobase million. A Log-rank test, the Mantel-Cox test, was used to determine *p* values. HR, hazard ratio.

Traditionally, prostate tumors have been graded on the Gleason scale (51–53). In this microscopic evaluation, the two most dominant patterns in the tumor biopsy are graded relative to normal prostate tissue, with the sum representing the Gleason score. Scores can range from 2 to 10, with 10 representing the least differentiated tumor cells that are most distinct from normal prostate tissue cells and that generally signify the worst prognosis. In practice, Gleason scores for prostate cancer patients typically range from 6-10. More recently, the International Society of Urological Pathology has further refined the diagnostic system (54). We plotted BRG1 and BRM mRNA expression as a function of Gleason score (Fig. 4A-B) and show the data in a heatmap format as well (Fig. 4C). Average values are presented in Fig. 4D. The data clearly indicate no correlation between BRG1 mRNA expression and Gleason score. Not surprisingly, there is also no correlation with BRM mRNA expression. Thus BRG1 mRNA expression is a prognostic indicator of prostate cancer patient outcome that is independent of Gleason score.

**Figure 4:**
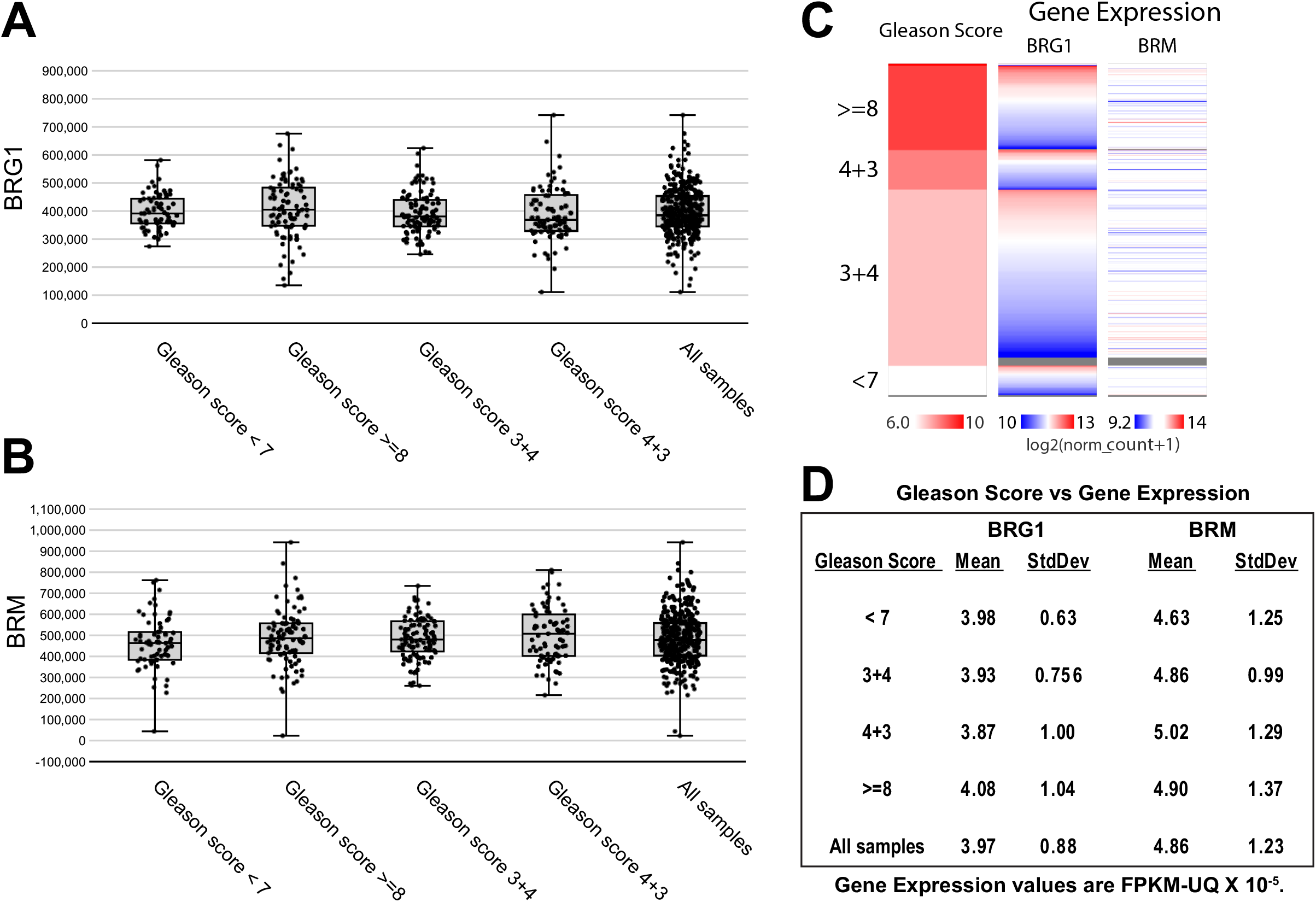
Neither BRG1 nor BRM mRNA levels correlate with prostate tumor Gleason score. **(A)** the ranges of BRG1 or **(B)** BRM mRNA levels from prostate tumor expression data in TCGA organized by the Gleason score for each tumor. The Project Betastasis website was used for data analysis and figure generation. FPKM, Fragments Per Kilobase of transcript per Million mapped reads, upper quartile normalized. **(C)** Heat map of BRG1 and BRM mRNA expression in prostate tumor samples with Gleason scores of <7, 7, or >7. **(D)** Mean BRG1 and BRM mRNA expression in prostate tumor samples with the indicated Gleason scores. SD, standard deviation.

### A BRG1 inhibitor diminishes prostate cancer cell survival in culture and in xenografts

The data suggest that targeting BRG1 may be of therapeutic benefit for prostate cancer. ADAADi is a biologic preparation isolated as a byproduct of the bacterial aminoglycoside-3’-phosphotransferase APH (3’)-III enzyme reaction (40,41). It demonstrates preference for targeting BRG1 over BRM in cell culture experiments (39) and phenocopies BRG1 knockdown in inhibiting lipid synthesis and in blocking drug-induced activation of ABC transporter gene expression in breast cancer cells (39,43). We therefore asked whether ADAADi might be used to inhibit prostate cancer cell proliferation and/or survival.

The PC3 prostate cancer cell line was derived from a patient’s metastatic prostate adenocarcinoma and is capable of anchorage independent growth in culture and of generating tumors in athymic nude mice (55). Treatment of PC3 cells proliferating in culture with increasing concentrations of ADDADi gave a dose-dependent decrease in cell viability (Fig. 5A). Annexin V staining indicated that the observed cell death in the presence of a sub-lethal concentration of ADAADi was due to apoptosis (Fig. 5B).

**Figure 5:**
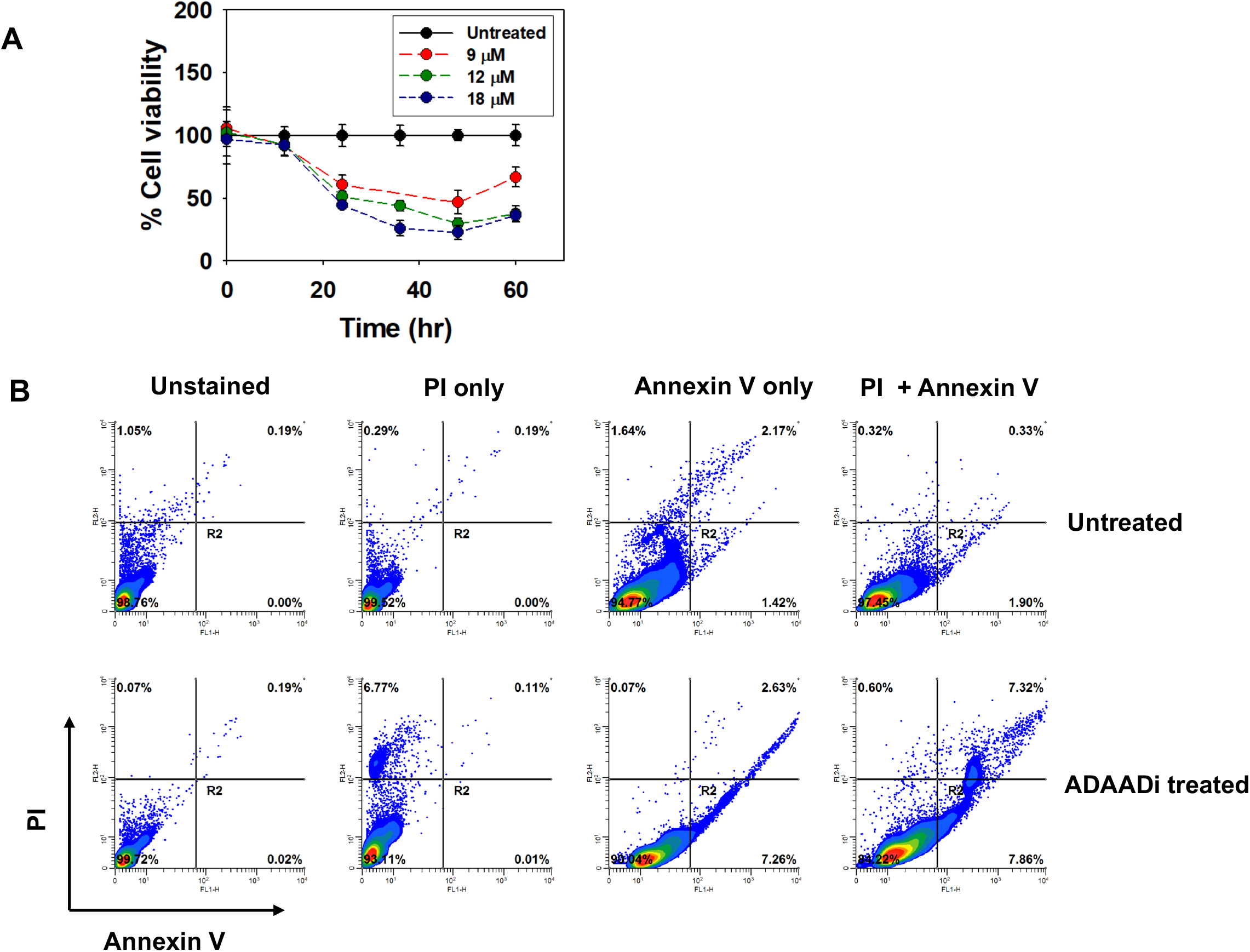
**(A)** ADAADi treatment inhibits PC3 cell proliferation. Cell viability was determined by MTT assay after 48 h in the presence of the indicated concentration of inhibitor. **(B)** ADAADi treatment of PC3 cells increases the frequency of apoptosis. Cells were treated with 5 uM ADAADi for 48 h and stained as indicated prior to FACS analysis.

Next, PC3 cells were injected into the flanks of athymic mice and subcutaneous tumor growth was monitored. ADAADi or PBS was injected directly into the subcutaneous tumor when the tumor reached a size of 200 mm^3^. In two independent experiments, every-other-day injections were executed for two weeks (Figs. 6A-B and 6C-D). A third experiment used the every-other-day protocol for two weeks and, after a one week break from treatment, an additional set of injections were performed, again, every other day for two weeks (Figs. 6E-F). The latter experiment also included direct injection of the parent aminoglycoside (kanamycin) as a control in addition to PBS (Fig. 6E). Tumor size was monitored during and after treatment.

**Figure 6:**
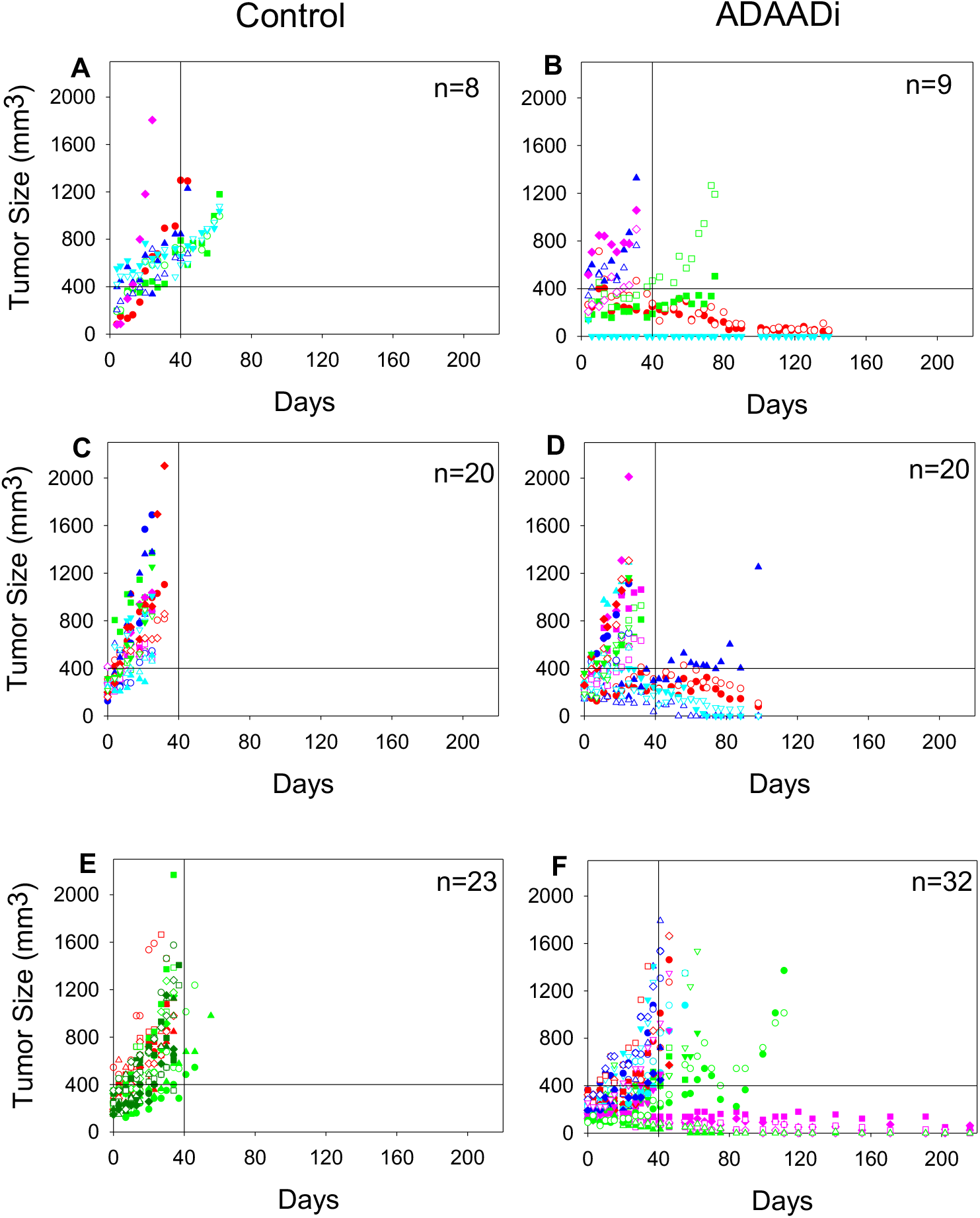
ADAADi injection into PC3-induced mouse xenografts reduces, and in some cases, inhibits tumor growth. Three independent xenograft experiments were initiated by injection of PC3 cells. Tumor size was plotted as a function of time following injection of ADAADi or the control solution directly into the tumors. The number of xenografts (n) in each cohort is indicated. Data points for each individual mouse are indicated by specific shapes and colors. In Experiment 1 **(A-B)** and Experiment 2 **(C-D)**, tumors were injected with PBS or ADAADi every other day for two weeks. In Experiment 3 **(E-F)**, tumors were injected with ADAADi or with kanamycin instead of PBS as a control every other day for two weeks and after a one week break from treatment, an additional set of injections were performed every other day for two weeks.

For clarity of presentation, tumor size data is presented for each of the three separate experiments that were performed (Figs. 6A-F). Mice injected with PBS or kanamycin showed consistently increasing tumor size and were euthanized when the maximum allowable tumor burden was observed, in accordance with IACUC protocols (Figs. 6A, C, E). ADDADi injected tumors showed a range of results. Some injected tumors expanded with similar kinetics to the control tumors (Figs. 6B, D, F). Others showed delayed expansion but nevertheless expanded to the point where the animal was sacrificed (e.g. - blue solid triangle in Experiment #2, green open and solid circles in Experiment #3). In contrast, some tumors failed to expand post ADAADi-injection, and a subset of these tumors completely dissipated (e.g. - red open and solid circles in Experiment #1). We conclude that ADAADi inhibited tumor growth in a subset of the treated animals. We also plotted survival data (Fig. 7), which showed a clear survival advantage for the treated animals and highlighted the fact that approximately 30% of the treated animals survived for up to 7.5 months without evidence of tumor or any observable physical or behavioral anomaly, at which time these remaining animals were euthanized. Necropsy of these “survivors” revealed no tumors in other tissues, with the kidneys, lungs, lymph nodes proximal and distal to the injection site, or spleen specifically examined in each individual. The data indicate that direct injection of ADAADi into the xenografts resulted in tumor growth inhibition and a survival benefit, suggesting that this inhibitor has potential as a therapeutic agent.

**Figure 7:**
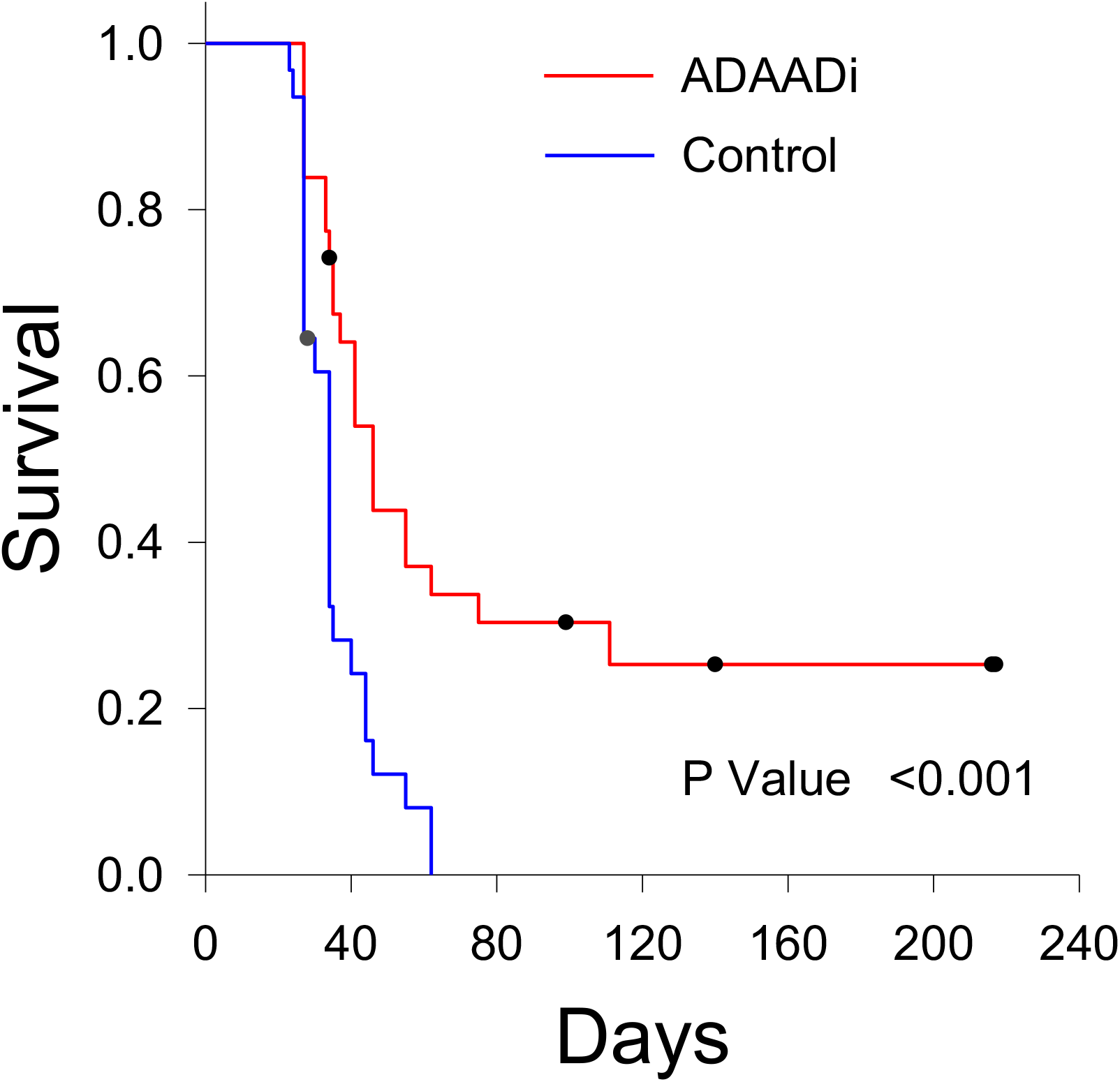
ADAADi injection into PC3-induced xenografts extends survival. Kaplan-Meier plots showing survival curves for all mice in the xenograft experiments presented in Fig. 6. A Logrank test was used for statistical analysis.

## DISCUSSION

The ATPases that are the catalytic subunits of human SWI/SNF enzymes have been linked to many types of cancer, but their functional contributions to oncogenesis is context dependent. Here we probed the TCGA database for expression and mutation frequency of BRG1 and BRM in prostate cancer. The analyses indicated elevated BRG1 and BRM mRNA and elevated BRG1 protein expression in prostate tumors relative to normal tissue. This is consistent with prior IHC studies of patient tumors (48). Although the TCGA data set did not contain information on BRM protein levels, a prior report revealed that BRM protein levels did not correlate with increased BRG1 protein in patient samples (49). A recent survey of multiple tumor types indicated that BRG1 expression was elevated in tumors whereas BRM was not (56). Mechanistically, it appears that there is a direct correlation between BRG1 mRNA and protein expression, while post-transcriptional regulation likely plays a role in determining overall BRM protein expression.

Elevated expression of a regulatory protein can suggest tumor promoting activity but could instead indicate tumor suppressive activity if there is elevated expression of a mutant protein. To address this question, we examined the 491 available sequences in the TCGA prostate cancer database for mutations in BRG1 and BRM. Neither BRG1 nor BRM are frequently mutated in the patient tumors. This result is consistent with and extends a prior report finding no mutations in BRG1 coding sequences from 21 patient tumors (57). Of the mutations identified in the TCGA database, a G775D alteration in BRG1 indicates a mutation in motif I of the conserved Snf2 ATPase domain (58). This residue is not conserved amongst the Snf2 family proteins. Nevertheless, we would predict that this mutation would render the protein inactive based on our prior structure-function studies (59). The I1214L mutation identified in BRG1 is outside motif VI of the ATPase domain, but the residue is conserved in many closely related Snf2 family members, and analysis of the *S. solfataricus* SWI2/SNF2 ATPase core complexed with DNA (60) suggests that this mutation likely impairs ATPase activity. The BRM R1159Q mutation identified in one patient lies in motif VI and is conserved across the entire Snf2 ATPase family. Mutation of this residue in the SMARCAL1 ATPase resulted in loss of ATPase activity (61). The other identified mutants are outside the conserved SNF2 ATPase domain, and the potential functional consequences are unknown.

Stratification of BRG1 expression amongst prostate tumor samples revealed that elevated BRG1 mRNA expression correlated with poor patient outcome. BRG1 expression is therefore a new marker for prostate cancer survival. This finding provides additional evidence that elevated BRG1 expression, not BRG1 mutation, is a marker for poor prognosis in an increasing number of cancer types. Prior work has demonstrated that high BRG1 mRNA expression inversely correlates with patient survival in breast cancer (27,28,39), colorectal cancer (62,63), and neuroblastoma (29). In addition, BRG1 is required for various aspects of cancer cell survival, proliferation, and function in HeLa cells (64), leukemia cells (65,66), breast cancer(33,43), hepatocarcinoma (67), colorectal cancer (68), neuroblastoma (29), melanoma (30,69,70), and certain medullablastoma tumors (71). Recent evidence indicates that the fusion between the TMPRSS2 gene and the ETS family transcription factor, ERG, that occurs in half of prostate cancers, mediates its oncogenic effect at least in part by interacting with the human SWI/SNF enzymes and re-directing its chromatin interactions across the genome (72). Thus the chromatin remodeling enzyme and presumably its enzymatic function is a required component contributing to prostate oncogenesis. Collectively, the data contrast with the idea of BRG1 acting as a tumor suppressor in all cancer types, support the idea of context dependent function of SWI/SNF ATPases in cancer, and raise the idea that BRG1 and/or BRM can, in some contexts, be drivers of oncogenesis (16).

We compared BRG1 expression to Gleason score, the commonly used staging system for prostate tumors (52), and found no correlation. The conclusion was based on analysis of 568 patient samples in the TCGA prostate cancer database. Our results indicate that BRG1 expression and Gleason score are therefore independent prognostic indicators of patient outcome. This point has been debated in the past, with one prior report finding correlation (49) and another finding no correlation (48). The smaller size of the respective sample pools in these studies (46 and 64 patient samples, respectively) may have contributed to the differing results.

In this report, we demonstrate that an inhibitor that shows specificity for BRG1 can be an effective tool in inhibiting prostate cancer cell survival in tissue culture and in xenografted prostate tumors. Prior studies have demonstrated that ADAADi is effective against numerous cancer cell types in culture (39,40), but this is the first demonstration that it is effective in an animal tumor model. The work therefore extends the proof-of-principle that targeting a chromatin remodeling enzyme can be an effective strategy for cancer treatment.

Specifically targeting the BRG1 enzyme as a therapeutic approach for cancer is an emerging idea. There are always drawbacks to targeting an essential protein or biological process, but we submit that differential effectiveness in cancerous versus normal cells will ultimately dictate the success of this strategy. Differential effectiveness is the reason that classical chemotherapy drugs that target cancers based on increased rate of cell division in tumor versus normal cells have been utilized for decades. So, despite BRG1 being ubiquitously expressed and functional in normal cells, there can be, and in fact are, cancer-specific roles for BRG1. In triple negative breast cancer, BRG1 is specifically required for the upregulation of lipid and fatty acid synthesis enzymes that are required to produce elevated levels of these building blocks for rapid cell division; knockdown or inhibition of BRG1 reduced overall de novo lipid synthesis in the cancer cells but not in breast epithelial cells (43). Another example of a cancer-specific role for BRG1 comes for other studies in triple negative breast cancer cells where it was shown that BRG1 mediates the induction of ABC transporters in response to treatment of cells with chemotherapeutic drugs. Knockdown or inhibition of BRG1 prevented induction of ABC transporters and resulted in increased intra-cellular retention of the drugs and increased chemosensitivity, raising the possibility that a BRG1 inhibitor could be an effective adjuvant treatment to classical chemotherapy drugs (39). The data support the idea that inhibition or reduction of BRG1 could preferentially impair cancer cell growth and function relative to normal cells. Delivery, like with many inhibitory drugs, will be an issue that will require further development. However, it is apparent based on the work presented here that direct delivery of the inhibitor to the tumor can be effective in tumor reduction without impairing lifespan or apparent health of the treated individual. Presumably this means that any effects of the treatment on the functions of BRG1 in normal cells was minimal or non-existent.

Finally, the work will be greatly advanced once the inhibitory molecule(s) present in the ADAADi preparation are identified. Work in our labs continues to address this problem. In addition to better defining ADAADi, our work indicates that screens of chemical libraries should be performed to identify novel inhibitors of BRG1, which we would predict would have immediate pre-clinical relevance for future therapeutic approaches to cancers where BRG1 expression is elevated relative to normal tissue.

## CONCLUSIONS

BRG1, one of the two SNF2 family ATPases that act as the mutually exclusive catalytic subunit of human SWI/SF chromatin remodeling enzymes, showed elevated expression in primary prostate tumors relative to normal cells. Mutation or amplification of BRG1 was observed in <1% of tumors. BRG1 expression was inversely correlated with patient survival and did not correspond to Gleason score, indicating it is an independent prognostic indicator for patient survival. An inhibitor of BRG1 inhibited prostate cancer cell line survival in culture, reduced tumor growth and extended survival in xenografted mouse tumors. The data support the idea that targeting BRG1 is an effective anti-tumor strategy.

## ACKNOWLEDGEMENTS

This work was partially supported by NIH grants GM56244 to ANI and EB014869 to JAN. RR was supported by UGC non-net fellowship. RM also acknowledges funding from DST-PURSE (PAC-JNU-DST-PURSE-462 (phase II)). We wish to thank Peter Pryciak for discussion and Hanna Witwicka for technical assistance and discussion.

